# Sequence determinants of the aggregation of proteins within condensates generated by liquid-liquid phase separation

**DOI:** 10.1101/2020.12.07.414409

**Authors:** Michele Vendruscolo, Monika Fuxreiter

## Abstract

The transition between the native and amyloid states of proteins can proceed via a deposition pathway via oligomeric intermediates or via a condensation pathway via liquid droplet intermediates generated through liquid-liquid phase separation. Here we investigate the sequence determinants of aggregation from within the droplet state based on generic interactions. We describe a model in which these sequence determinants can be captured by three features, the droplet-promoting propensity, the aggregation-promoting propensity and the binding mode entropy. By using this approach, we propose a formula to identify aggregation-promoting mutations in droplet-forming proteins. This analysis provides insights into the amino acid code for the conversion of proteins between liquid-like and solid-like condensates.

## Introduction

Recent advances have revealed that proteins exhibit a complex phase behaviour, as they can populate the native, droplet and amyloid states [1, 2]. It has also become clear that the amyloid state can be reached either through a deposition pathway [3, 4], or through a condensation pathway, going through a liquid-liquid phase separation (**Figure 1**) [5–7]. The discovery of the phenomenon of liquid-liquid phase separation is highlighting a variety of previously unknown biological processes [8, 9]. Since the transition between the native and droplet states is usually reversible under cellular conditions, as opposed to that between the native and amyloid states, the droplet state can be rather safely exploited for functional processes in living systems [10, 11].

**Figure 1.**
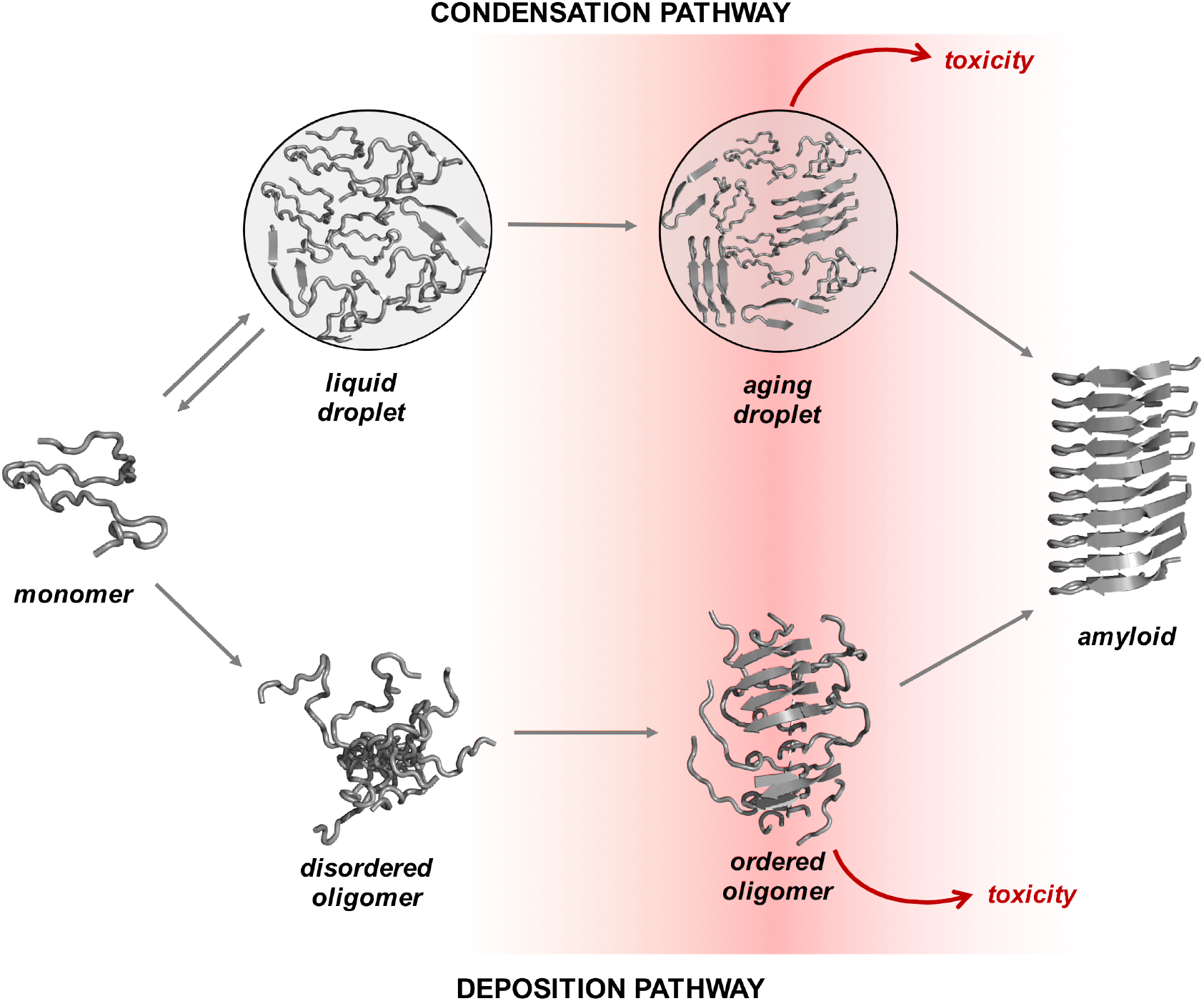
Overview of the condensation and the deposition pathways for amyloid formation. Along the deposition pathway, proteins move from the native state to the amyloid state through increasingly ordered oligomer aggregates [3, 4]. Some of these oligomers are highly cytotoxic [67, 68] according to an amino acid code that has been partly elucidated [69]. Along the condensation pathway, proteins that undergo condensation convert from the native state to the amyloid state through a dense liquid state (the droplet state). The stability of these different states, as well as the conversion rates between them, are different for different proteins. For many proteins under cellular conditions, the native and droplet states are metastable, and the conversion rates between the droplet and fibrillar states are fast. For certain proteins, the droplet state is functional, and it is stabilized by extrinsic factors, such as RNA and post-translational modifications. The maturation of droplets into the amyloid state creates cytotoxic intermediates [6, 13, 15, 16].

Upon dysregulation of the native-to-droplet transition, however, the droplet state can evolve into a transient gel form, which can then further proceed to the amyloid state (**Figure 1**) [12]. This gel state may give rise to cytotoxicity, as it can less readily revert to the native state [6, 13–16]. In particular, proteins can remain entangled in the gel form because they recruit other cellular components [17]. These observations prompt the question of whether there is an amino acid code determining the condensation pathway of amyloid formation [18, 19], in analogy to the amino acid code that has been identified for the deposition pathway [20, 21].

Structural studies indicate that characteristic signatures of the amyloid state, such as cross-β elements are also present in the droplet state [22–24]. Furthermore, labile β structures are enriched in droplet-forming sequences [25]. Here we focus on how the amino acid sequences encode distinct physico-chemical properties that drive the conversion between liquid-like and solid-like condensates.

We compare the nature of the interactions along the condensation pathway between droplet-promoting regions and amyloid-promoting regions. We propose a model in which the condensation pathway is driven by conformationally heterogeneous elements, which have been referred to as labile structures [25, 26]. Our analysis indicates the possibility of predicting aggregation-promoting mutations in droplet-forming proteins based on generic interactions and the ability of sequence elements to switch between different interactions modes. We thus offer insights into the mechanisms and sequence determinants governing the condensation pathway between liquid-like and solid-like condensates and the modulation of this process by pathological mutations.

## Results

### A framework for the generic approach

Formation of condensed states, either the liquid-like droplet state or the solid-like amyloid state, is generic to proteins [3, 27]. Both states are stabilised by generic interactions, in accord with the wide variety of sequence motifs identified by experimental studies capable of driving liquid-liquid phase separation [27]. Here, we hypothesized that the condensation pathway can be described by sequence-encoded biophysical properties, and aimed at analysing these generic interaction behaviors.

### Datasets of protein condensates representing the droplet and amyloid states

We assembled a database of droplet-forming proteins (DROP), termed ‘droplet-drivers’ (DROP-D), using three databases: PhaSepDB (http://db.phasep.pro) [28], PhaSePro (https://phasepro.elte.hu) [29], and LLPSDB (http://bio-comp.org.cn/llpsdb) [30] (Methods, **Table S1**). Droplet-client proteins (DROP-C), which require an interacting partner for phase separation, are derived from LLPSDB database [31]. Regions of droplet-driver or droplet-client proteins that mediate droplet formation are assembled from the PhaSePro dataset [29] (Methods, **Table S1**). Droplet-promoting regions (DPRs) are defined based on an evidence for spontaneous phase separation (Methods, **Table S1**). In addition, low-complexity aromatic-rich kinked segments (LARKs) are used to represent structural hallmarks of droplet formation [25].

The amyloid state is populated by both pathological and functional amyloids (AMYLOID), which are derived from the AmyPro database (http://amypro.net) [32] (Methods, **Table S1**). Amyloid-promoting regions (APRs) are derived from the AmyPro database [32], which include amyloid core-forming cross-β structures and their flanking regions (Methods, **Table S1**).

In addition to the above categories, which represent proteins forming liquid-like and solid-like condensed states [33], we assembled proteins associated with both states. Proteins with an evidence for both droplet proteins and amyloid formation are defined as ‘droplet-amyloids’ (DROP-AMYLOID, Methods, **Table S1**). This category contains proteins with prion-like domains (e.g. Sup35) and proteins, which mature into fibrillar aggregates (e.g. tau). Proteins associated with inheritable traits in *Saccharomyces cerevisiae* [34], are defined as yeast prions (YPRIO, **Table S1**). These proteins represent functional amyloids, but we listed them separately from the droplet and amyloid datasets (Methods, **Table S1**).

The assembled datasets exhibit a wide range of solubility values [35] (**Figure 2A**).

**Figure 2.**
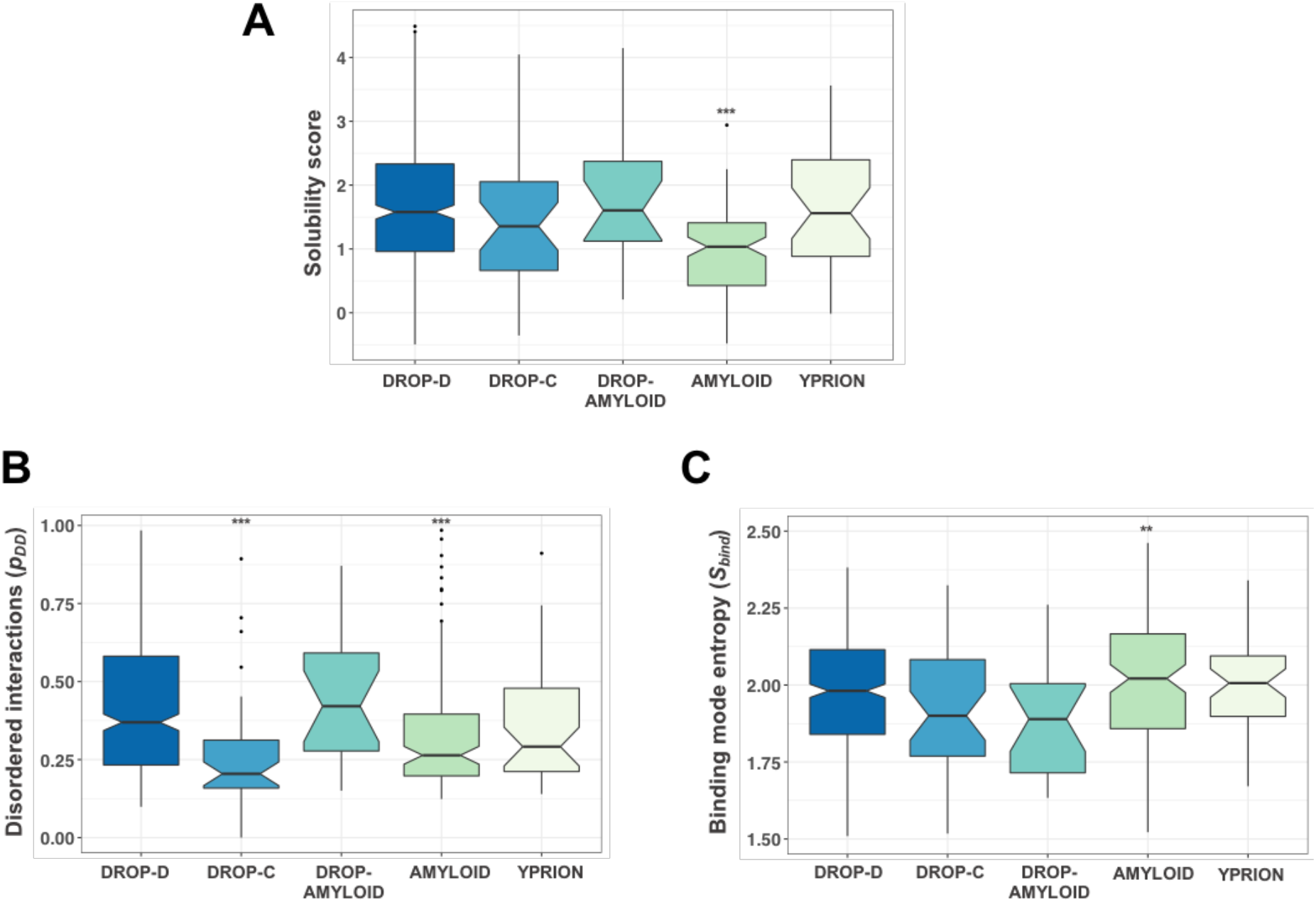
Comparison of solubility and interactions of amyloid-forming and droplet-forming proteins. **(A) Comparison of the solubility scores of different datasets of droplet and amyloid proteins.** Droplet-driver proteins (DROP-D, blue, **Table S1**), which spontaneously undergo liquid-liquid phase separation [27] and droplet-amyloid proteins (DROP-AMYLOID, aquamarine, **Table S1**), which exhibit both droplet and amyloid behaviors, have the highest solubility scores by CamSol [35], while amyloid-forming proteins (AMYLOID, green) in the AmyPro database (http://amypro.net) [32] have the lowest solubility scores. Droplet-client proteins (DROP-C, light blue, **Table S1**), which require partner-interactions for droplet formation, and yeast prions (YPRION, light green), with inheritable traits in *Saccharomyces cerevisiae* [34], have intermediate solubility scores. Statistical significances by Mann-Whitney test (***p< 10^−3^) show that amyloid-forming proteins have significantly lower solubility scores than droplet-forming proteins, while other classes sample a continuum between these two extremes. **(B) Comparison of the disordered binding modes (*p_DD_*) of droplet and amyloid proteins.** Droplet-driver and amyloid-forming proteins exhibit distinct interaction behaviors. FuzPred [42] predicts disordered binding modes (Methods, Eq. 1) for droplet-driver proteins, and ordered binding modes for amyloid proteins. Droplet-amyloid proteins have disordered interaction behaviors as droplet-drivers, while yeast amyloids and droplet-clients have ordered binding modes as amyloids. Statistical significances by Mann-Whitney test (***p< 10^−3^) show that amyloid-forming proteins and droplet-client proteins exhibit significantly more ordered interactions than droplet-forming proteins, while yeast prions and droplet amyloids sample both interaction behaviors. **(C) Comparison of the binding mode entropy** (*S_bind_*) **of droplet-forming and amyloid-forming proteins.** Amyloid-forming proteins exhibit high binding mode entropy, as they can sample both disordered and ordered interactions. Droplet and droplet-amyloid proteins have a strong preference for disordered binding (low S_bind_). S_bind_ values were computed by FuzPred [41] (Methods, Eq. 3). Statistical significances by Mann-Whitney test (** p < 10^-2^) show that amyloid-forming proteins are more prone to multimodal interactions than droplet-forming proteins, while the other classes sample a similar range of binding entropies. Statistical significances as compared to droplet driver proteins were computed by Mann-Whitney test using the R program.

### Comparison of the binding modes of amyloid-forming and droplet-forming proteins

Increasing evidence demonstrates that proteins can exhibit a wide range of binding modes by sampling a continuum between ordered and disordered bound states [36, 37], and may switch between such states by post-translational modifications or according to the cellular context [38]. An ordered binding mode is mediated by specific contact patterns [39], which create a free energy landscape with a well-defined minimum corresponding to the bound state. By contrast, a disordered binding mode involves multimodal interactions [40], which create a heterogeneous ensemble of configurations with multiple minima in the free energy landscape of the bound state. We previously demonstrated that both ordered and disordered binding modes of proteins can be predicted from their amino acid sequences [41, 42]. We also showed that the propensity of proteins to spontaneously undergo liquid-liquid phase separation depends on the probability of being disordered in their free states (*p_D_*) and the probability of remaining disordered upon binding (*p_DD_*) [27].

In this approach, droplet-driver proteins tend to be disordered in both their free and bound forms [27], while amyloid-forming proteins tend to be ordered in both forms (**Figures 2B** and **S1A**). In addition, droplet-clients tend to resemble the ordered interactions of amyloid-forming proteins, while yeast prions, and in particular droplet-amyloid proteins exhibit disordered interactions, which do not deviate significantly from droplet-driver proteins (**Figures 2B** and **S1A**).

Amyloid-forming proteins exhibit the ability to switch between different binding modes, therefore having a high binding mode entropy (*S_bind_*, Eq. 3) (**Figure 2C**). By contrast, droplet-driver and droplet-client proteins, and in particular droplet-amyloid proteins, exhibit low binding mode entropy (**Figure 2C**), reflecting a strong preference for forming disordered interactions in the condensates. Interestingly, the predictions indicate that yeast prions are less prone to switch from disordered to ordered interaction modes than amyloid-forming proteins.

Taken together, this analysis identifies different binding modes in droplet-forming and amyloid-forming proteins, the latter being associated with a greater propensity to switch between binding modes.

### Droplet-promoting and amyloid-promoting regions can populate both the droplet and the amyloid states

Droplet-promoting regions are enriched in disorder-promoting residues (serine, glycine and proline) [43] as compared to amyloid-promoting regions, while depleted in order-promoting residues (valine and leucine) (**Figure S2**). This is in accord with previous observations that proline and glycine composition controls the properties of elastomeric and amyloid-like fibrils [44].

The two extreme scenarios leading to the condensation pathway are represented by low complexity regions (LARKs) [25], which have high droplet-promoting propensity (*p_DP_*) [27] and low amyloid-promoting propensity (*p_AP_*) [35], and by amyloid cores, which have low droplet-promoting propensity (**Figure S3D**) and high amyloid-promoting propensity (**Figure S3C**). These structural elements exhibit contrasting binding modes (*p_DD_*) (**Figure 3A**) and binding mode entropy (*S_bind_*) (**Figure S3B**). By contrast, droplet-promoting and amyloid-promoting regions do not exhibit a significant difference in these interaction behaviors (**Figures 3A** and **S3B**) neither in their amyloid-promoting (**Figure S3C**) and droplet-promoting propensities (**Figure S3D**). This analysis indicates that residues in amyloid-promoting regions (**Figure 3B**) and in droplet-promoting regions (**Figure 3C**) can populate both the droplet and the amyloid states. Indeed, residues in amyloid-promoting and droplet-promoting regions were predicted in both amyloid and droplet states (**Figures 3B, 3C**). This analysis is in accord with structural studies reporting amyloid-like labile β elements in the droplet state [22, 24].

**Figure 3.**
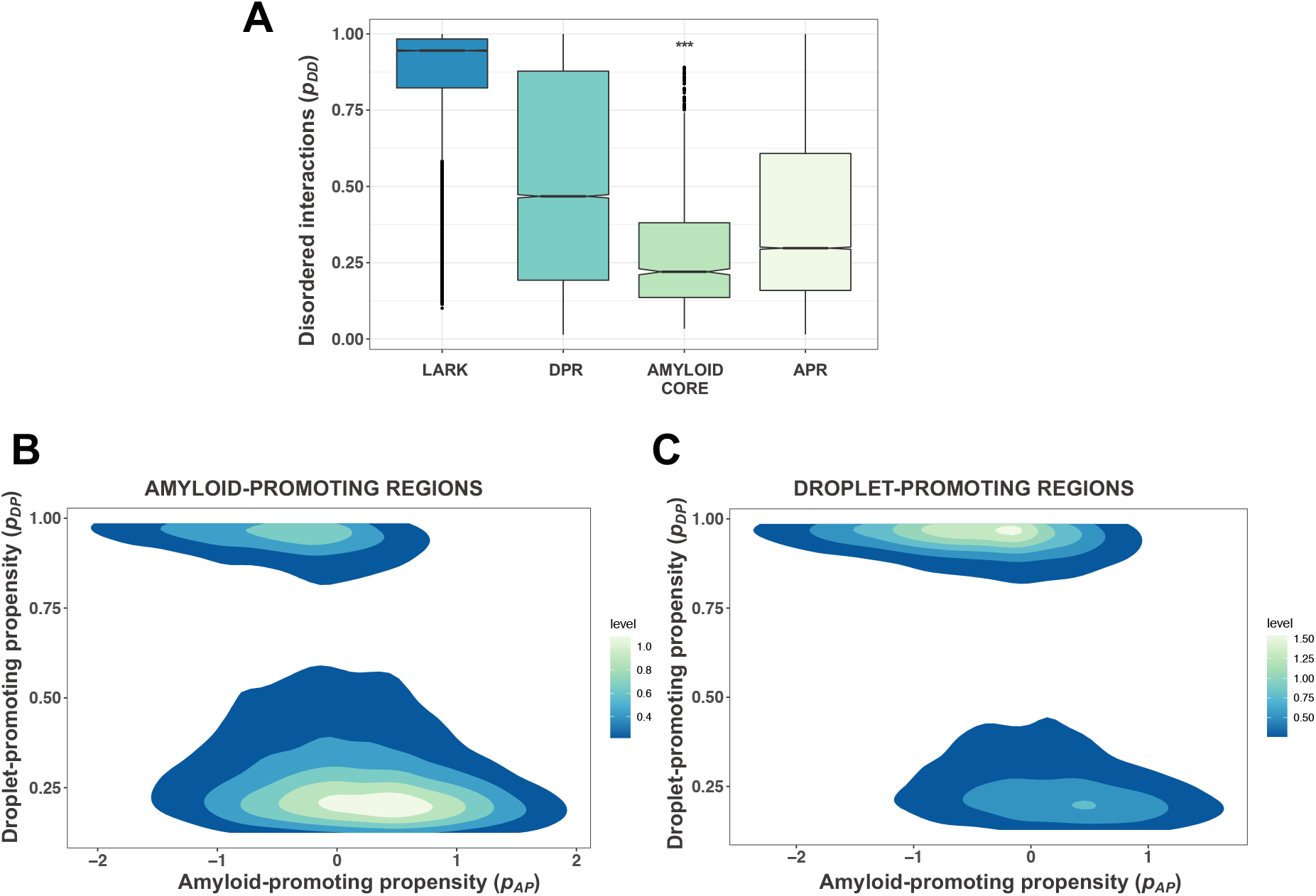
Experimentally-observed droplet-promoting and amyloid-promoting regions can sample both states. **(A) Comparison of the disordered binding modes (*p_DD_*) of droplet-promoting** (DPR, aquamarine) **and amyloid-promoting** (APR, light green) **regions.** Interactions of droplet-promoting and amyloid-promoting regions (**Table S1**) vary in wide range between these disordered and ordered binding modes, exemplified by LARKs [25] (blue) and amyloid cores [32] (green). The statistical significance in reference to droplet-promoting regions, as computed by Mann-Whitney test using the R program (*** p < 10^−3^), demonstrates that interaction properties of droplet-promoting and amyloid-promoting regions overlap. **(B,C) Comparison of the droplet-promoting propensity** (p_DP_, y-axis) **and amyloid-promoting propensity** (p_AP_, x-axis) **of amyloid-promoting (B) and droplet-promoting regions (C).** Droplet-promoting propensities were computed by FuzDrop [27] and amyloid-promoting propensities were derived from the solubility scores by CamSol [35]. The amyloid state is characterised by low p_DP_ and positive p_AP_ values, while the droplet state has high p_DP_ and negative p_AP_ values. Residues in amyloid-promoting regions can also populate the droplet state, and residues in droplet-promoting regions can also sample the amyloid state.

In addition, we found that about half of the amyloid-promoting regions contained short droplet-promoting regions (≥ 10 residues, **Table S2**), and that about half of the droplet-promoting regions contained short amyloid-promoting regions (≥ 6-residues, **Table S2**). The droplet-promoting propensity increased with the distance from the amyloid core to reach a maximum at ~25-35 residues from the core (**Figure S4A,C**), while the amyloid-promoting propensity did not depend considerably from the position of the flanking region (**Figures S4B,D**). Along these lines, we observed a widespread co-occurrence of droplet-promoting and amyloid-promoting regions in different proteomes (*Homo sapiens* 87%, *Saccharomyces cerevisiae* 88%, in *Caenorhabditis elegans* 89%, **Table S3**) based on predictions by the CamSol [35], Tango [20] [45] and FuzDrop methods [27].

Taken together, this computational analysis indicates that, although droplet-promoting regions and amyloid-promoting regions have distinct binding modes, these regions can sample both the droplet and the amyloid states.

### A droplet landscape illustrates the propensity for the droplet and amyloid states

The computational analysis reported above indicates that the conversion between the droplet and amyloid states requires a switch from disordered to ordered binding modes involving sequence regions that can sample both states. As illustrated by the case of the prion-like domain of TDP-43 (residues 262-414), residues that drive the condensation pathway can interact in both disordered and ordered binding modes (**Figure 4A**). However, the frequency of disordered interactions, which favor the formation of the droplet state, is higher than those of ordered modes, which promote the stabilization of the amyloid state (**Figure 4A**).

**Figure 4.**
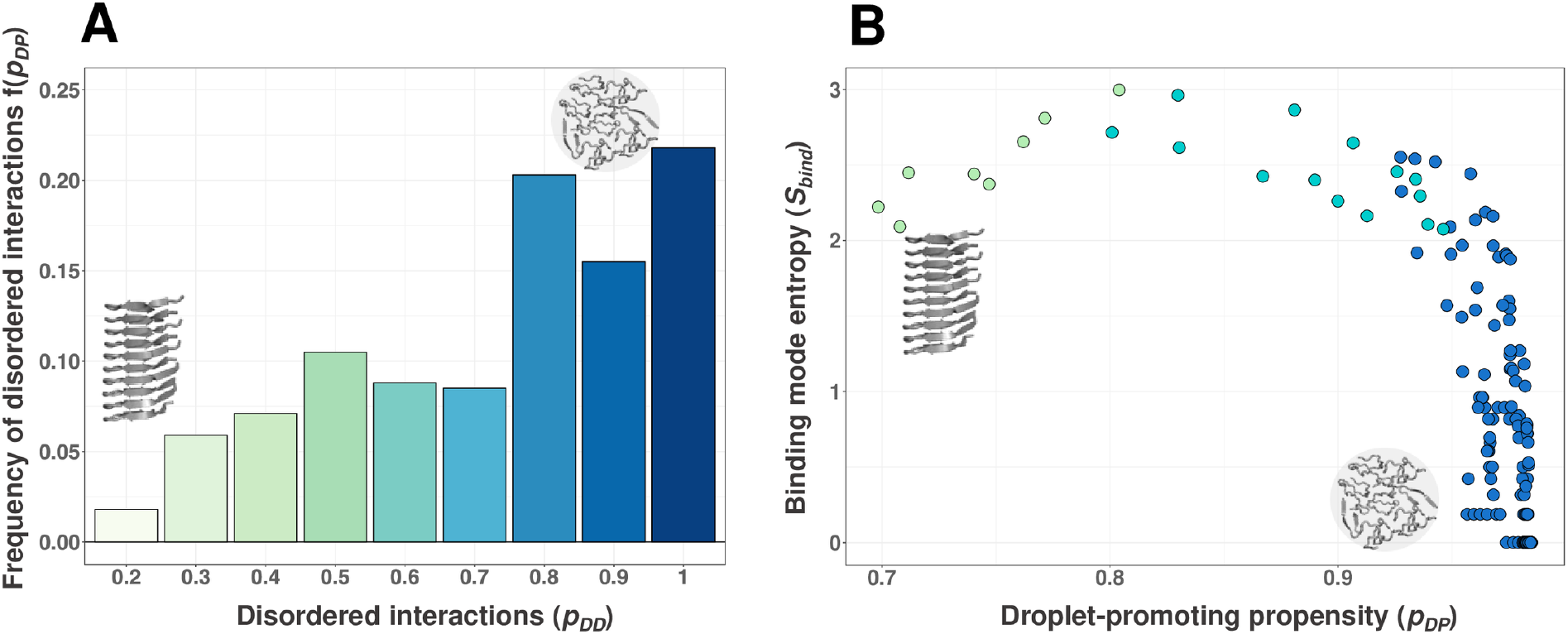
Droplet landscape of the prion-like domain of TDP-43. **(A) Frequency of different binding modes, f(p_DD_), for the residues in the aggregation hot-spot (residues 312-342) of TDP-43**. The high values of f(p_DD_) for large p_DD_ values indicate that the prion-like domain of TDP-43 preferentially samples disordered interactions corresponding to the droplet state (blue, right) [42], but also that it can form ordered interactions corresponding to the amyloid state (green, left). **(B) Droplet landscape of the TDP-43 prion-like domain (residues 262-414).** The droplet-promoting propensity (p_DP_) is shown on the x-axis [27], and the binding mode entropy (S_bind_) is shown on the y axis; S_bind_ is computed for each residue from the frequencies displayed in panel **(A)** (Eq. 3) [41]. Residues that tend to remain in the droplet state are found in the bottom right regions (blue), since they have high p_DP_ and low S_bind_ values, reflecting a strong preference for disordered interactions in the droplet state. Residues that are likely to convert to the amyloid stat, such as the amyloid-core of TDP-43 (321-330 residues, light green), are found in the upper left region (green), since they have low p_DP_ and high S_bind_ values, reflecting a tendency to change binding modes. Residues in the hot-spot region (aquamarine) have high p_DP_ and S_bind_ values, and therefore can facilitate the switch between disordered and ordered binding modes.

Based on this analysis, we propose to characterize conversion from the droplet state to the amyloid state using a droplet landscape, which is defined by the droplet-promoting propensity (*p_DP_*) of the residues and their binding mode entropy (*S_bind_*). Residues that are likely remaining in the droplet state are found in the bottom right region of the droplet landscape, since they have high droplet-promoting propensities and low binding mode entropy (**Figure 4B**). By contrast, residues with high propensity to form the amyloid state are found in the top left region of the droplet landscape, since they have low droplet-promoting propensity and high binding mode entropy (**Figure 4B**) [46]. Most residues of the TDP-43 aggregation hot-spot region (residues 312-342) are located in the top right region of the droplet landscape, indicating that these residues can exhibit both ordered and disordered binding modes, and can therefore facilitate the conversion from the droplet to the amyloid state (**Figure 4B**) [47].

Overall, the droplet landscape quantifies the contribution of different residues to the different components driving the condensation pathway (**Figure 4B**). An analysis of the droplet landscape suggests that residues with a wide range of binding modes can drive the conversion from the droplet state to the amyloid state, while those with strong preference for disordered binding modes tend to remain in the droplet state (**Figure S5**).

### Impact of ALS-associated mutations on the condensation pathway

We used these insights to characterise ALS-associated mutations shown to promote aggregation of liquid droplets (**Table S4**): FUS G156E [6], G187S, G225V, G230C, G399V [48], P525L [15]; hnRNPA1 D262V [5], D214V [49]; hnRNPA2 D290V [50]; tau P301L, P301S, A152T [51]; TDP-43 A321V [47], G298S, M337V [52], A315T [53]; TIA1 P262L, A381T, E384K [54]; UBQL2 P506T [55]. As compared to the wild-type proteins, ALS-associated mutations shifted the droplet landscape towards lower droplet-promoting propensity (p_DP_) and higher binding mode entropy (S_bind_) (**Figure S6**). Thus, we expressed the change in aggregation propensity within a droplet, 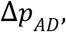, as a function of Δ*p_AP_*, Δ*p_DP_*, Δ*S_bind_* (see Methods), where Δ*p_AP_* = Δ*p_AP_*(*A_m_*) − Δ*p_AP_*(*A_wt_*) is the difference in amyloid-promoting propensities between the mutated (A_m_) and wild-type (A_wt_) residues, Δ*p_DP_* = Δ*p_DP_*(*A_m_*) − Δ*p_DP_*(*A_wt_*) is the corresponding difference in droplet-promoting propensities, and Δ*S_bind_* = *S_bind_*(*A_m_*) − *S_bind_*(*A_wt_*) is the corresponding difference in binding mode entropy (Methods). Using recovery rates of non-ALS associated FUS mutants (G154E, G156D, G156Q, G156P) and the ALS mutant G156E [56], we then trained a support vector model (SVM) for Δ*p _AD_* (see Methods) (**Figure 5A**). We note that some of these mutations, for example G156E, decrease the amyloid-promoting propensity Δ*p_AP_*, while considerably increasing the binding mode entropy Δ*S_bind_* as compared to the wild-type protein. The non-ALS associated G154E mutation has a considerably lower binding mode entropy than G156E and does not promote FUS aggregation. We then applied this model to predict the impact of the ALS-associated mutations on the aggregation propensity within droplets. For all the ALS-associated mutations we found high predicted aggregation propensity as compared to the non-ALS associated FUS mutations, which were experimentally shown not to promote aggregation (**Figure 5B**) [56].

**Figure 5.**
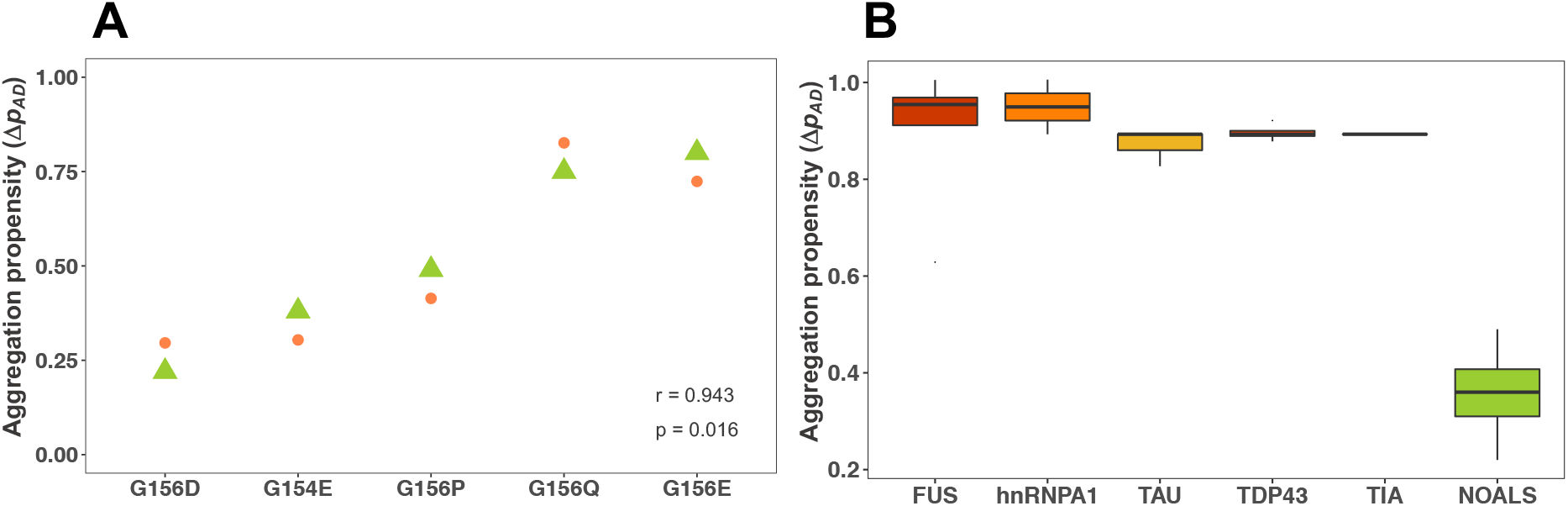
Prediction of the change of aggregation propensity within droplets (Δp_AD_) for ALS-associated mutations. **(A) Comparison of the change upon mutation of experimental (green) and predicted (orange) aggregation propensity within droplets of FUS mutants.** The experimental change in aggregation propensity was derived from reference [56]. The predicted change in aggregation propensity (Δp_AD_) was predicted using a SVM model with three features (change in droplet-promoting propensity, Δ*p_AD_*, change in aggregation-promoting propensity Δ*p_AD_*, and change binding mode entropy Δ*S_bind_*) that can be calculated from the sequence of a protein (Methods). **(B) ALS-associated mutations of FUS, hnRNPA1, tau, TDP-43, TIA1 have higher predicted change in propensity of aggregation within droplets than mutations not associated with ALS.** ALS-associated single mutations (orange) were shown to promote aggregation (FUS G156E [6], G187S, G225V, G230C, G399V [48], P525L [15]; hnRNPA1 D262V [5], D214V [49]; hnRNPA2 D290V [50]; tau P301L, P301S, A152T [51]; TDP-43 A321V [47], G298S, M337V [52], A315T [53]; TIA1 P262L, A381T, E384K [54]; UBQL2 P506T [55]) (**Table S4**) are predicted to induce large changes in propensity of aggregation within droplets than non-ALS associated FUS mutations G145E, G156D, G156P, G156A (green). Only proteins with multiple mutations are shown for clarity.

Thus, we propose that using three sequence-based features (droplet-promoting propensity, aggregation-promoting propensity and binding mode entropy) it is possible to predict the impact of a mutation on the condensation pathway. Systematic studies on aggregation-promoting mutations under consistent experimental conditions would be required for robust parametrisation. In particular more mutations, which do not induce amyloid formation will need to be analysed.

### Discussion and Conclusions

Most proteins are capable of converting into the amyloid state when their concentration exceeds their solubility limits [57], which can take place via disordered oligomers through the deposition pathway [3, 4]. It has been recently proposed that the droplet state is also widespread and likely to be accessible to most proteins [27]. Although the droplets formed through liquid-liquid phase separation can be functional, with new biological roles being discovered with rapid pace [8], they can also convert to amyloid state through the condensation pathway [5–7, 51].

The possibility for proteins of populating different states creates a challenge for the protein homeostasis system, since dysregulated transitions into dysfunctional assemblies can initiate pathological processes [3, 12–14]. It is therefore important to understand the sequence determinants driving liquid droplets towards solid-like aggregates. Our computational analysis indicates that that droplet-forming and amyloid-forming regions tend to exhibit different binding modes, while regions promoting both states tend to sample multiple binding modes. Along these lines, our study suggests that the condensation pathway is driven by residues that are capable of multiple binding modes, which are often associated with conformationally-heterogeneous structural elements [26, 47, 58].

We characterised the aggregation-promoting propensity of residues via the condensation pathway through a droplet landscape approach. As illustrated in the case of the prion-like domain of TDP-43, residues that tend to form droplets as they have high droplet-promoting propensity, and have high binding mode entropy are likely convert to amyloids, while those with strong preference for disordered interactions tend to remain in the droplet state. This analysis indicates that the structural ordering towards the amyloid state depends on the binding modes of the proteins in the condensates. In case of homotypic interactions (low binding mode entropy), the condensates will mature into the amyloid state, whereas in case of heterotypic (high binding mode entropy) interactions aggregation will be slowed down. The latter interactions, however, may also lead to promiscuity and cause cytotoxicity [59].

Indeed, most ALS-associated mutations experimentally shown to promote the condensation pathway also tend to increase multimodal interactions. This analysis may explain why amyloid-like structural elements exhibiting ordered interactions are also observed in droplets [22, 24], where disordered interactions dominate [60, 61]. We have also discussed how the aggregation propensity within droplets of a protein can be estimated from its amino acid sequence based on the droplet landscape and amyloid-promoting propensity. Based on this approach, we have proposed that ALS-associated mutations, including FUS G156E, do not necessarily reduce solubility, but expand interaction multimodality and facilitate the conversion towards ordered interactions. This behavior is in contrast to FUS G154E, which is less prone to increase interaction multimodality and does not promote aggregation.

Recent experimental results indicate that biophysical properties of droplets are evolved for optimal physiological function, while disease-causing mutations lead to aberrant condensates [62], and the normal condensate properties can be restored by unrelated droplet-promoting sequences [62]. Non-specific interactions between FUS ALS mutants and nuclear import receptors were also capable of restoring normal FUS condensate properties [48]. Although droplet-related pathologies are due to different molecular mechanisms [12], these observations indicate that familial mutations interfere with generic interactions [63]. This is consistent with earlier computational studies [27] and the analysis in this work indicates that the impact of a mutation on droplet aggregation can be estimated without specific sequence motifs.

Taken together, our computational analysis suggests an amino acid code based on three features (droplet-promoting propensity, aggregation-promoting propensity and binding mode entropy) that drive the transition of liquid-like condensates into amyloid deposits, and can be used to rationalise the impact of disease-related mutations on this transition.

## Methods

### Datasets of droplet-forming proteins

Spontaneously phase-separating proteins were assembled from three public databases (**Table S1**): the PhaSepDB Reviewed dataset based on curated literature search (http://db.phasep.pro) [28], the PhaSePro database with regions mediating phase separation identified (https://phasepro.elte.hu) [29], and the LLPSDB database (http://bio-comp.org.cn/llpsdb) [30] with experimental conditions specified for *in vitro* liquid-liquid phase separation. Merging these datasets resulted in 426 droplet-driver proteins, which can spontaneously phase separate (DROP-D, **Table S1**) [64]. Droplet-client proteins, which phase separate via interactions with other partners, were assembled from the LLPSDB dataset (http://bio-comp.org.cn/llpsdb) (DROP-C, **Table S1**). 20 proteins with evidence for both droplet and amyloid formation based on the AmyPro database (http://amypro.net) [32] were defined as droplet-amyloids (DROP-AMYLOID, **Table S1**).

Droplet-promoting regions were assembled from the PhaSePro dataset, and were grouped based on the evidence for spontaneous, or partner-assisted phase separation in the LLPSDB dataset (DPR, **Table S1**). Low-complexity aromatic-rich kinked segments (LARKs) were used as a reference for droplet-promoting regions [25]. The LARK dataset contained 6075 6-residue long segments in 400 proteins, as in the original reference.

### Datasets of amyloid-forming proteins

120 amyloid-forming proteins were assembled from the AmyPro database (http://amypro.net) [32] (AMYLOID, **Table S1**). This dataset contained only proteins with UniProt identifiers, with peptides excluded. Droplet-amyloid proteins were also eliminated from the amyloid dataset and were defined as a separate category (**Table S1**). Amyloid-forming proteins were annotated as pathological and functional based on the AmyPro database (**Table S1**). 46 yeast proteins with heritable traits in *Saccharomyces cerevisiae* were defined as yeast prions (YPRION, **Table S1**) [34]. Most yeast prions had Hsp104 or Hsp70 dependence [34]. Amyloid-promoting regions (APRs, **Table S1**) were defined based on the AmyPro database, including both the cross-β spine forming amyloid cores and the flanking regions. Amyloid-promoting regions were defined based on the AmyPro database, including both the cross-β spine forming amyloid cores and the flanking regions.

### Disorder-to-disorder binding profiles

The disorder-to-disorder binding profile, *p_DD_*, is estimated for each amino *A_i_* as [42]

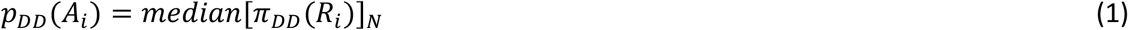

where π*_DD_*(*R_i_*) is the probability of disorder-to-disorder binding of *R_i_*, a region of 5-9 residues around *A_i_*, and N is the number of possible regions *R_i_*. We refer to π*_DD_*(*R_i_*) as the ‘binding mode probability’ of region *R_i_* because a value of 0 indicates a binding from fully disordered to fully ordered states, and a value of 1 indicates a binding from fully disordered to fully disordered states. *p_DD_* was computed using FuzPred [42].

### Binding mode entropy profile

The binding mode entropy profile (*S_bind_*) quantifies the variability of binding modes at the amino acid level. To define *S_bind_* we start by defining the frequency *f* of different possible binding modes for an amino acid *A_i_* as

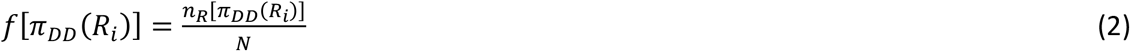

To calculate *f*, the binding modes are divided into discrete bins (usually 10), and *n_R_* is the number of binding regions within a given bin. The binding mode entropy profile (S_bind_) is then defined as the entropy of the frequencies of *f* [41]

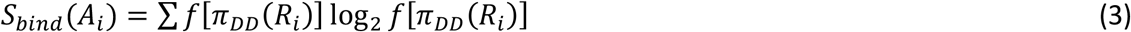

where the summation is over the [π*_DD_*(*R_i_*)] bins.

### Droplet-promoting propensity profiles

The droplet-promoting propensity profile p_DP_ quantifies the probability of spontaneous liquid-liquid phase separation [27]

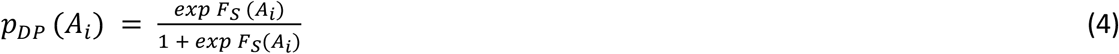

where *F _S_* (*A_i_*) is the scoring function for residue *A_i_*

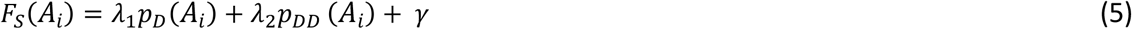

In this equation, *p_D_*(*A_i_*) is the probability for the disorder in the free state, and *p_D_*(*A_i_*) is the probability for disordered binding. *p_D_*(*A_i_*) approximates the conformational entropy in the unbound form, while *p_D_*(*A_i_*) estimates the conformational entropy of binding. λ_1_ and λ_2_ are the linear coefficients of the predictor variables and *γ* is a scalar constant (intercept), which were determined using the binary logistic model [27]. *p_D_* was derived from the disorder score as computed using the Espritz NMR algorithm [65]. The *p_DD_* values were predicted by FuzPred [42]. p_DP_ =0.60 is the threshold that a residue is involved in spontaneous phase separation.

### Amyloid-promoting propensity profiles

The amyloid-promoting propensity profile p_AP_ of a protein expresses its sequence-dependent probability to aggregate. Amyloid-promoting propensity profiles were obtained by the solubility profiles obtained by CamSol (p_CS_) [35]. p_AP_ = −p_CS_ and p_AP_ = 0.90 is the threshold, above which a protein likely aggregates [66]. The parameters *b* and *c* were determined by the R program.

### Prediction of the propensity of aggregation within droplets (Δp_AD_)

To calculate Δp_AD_ we expressed it as a function of sequence-based features

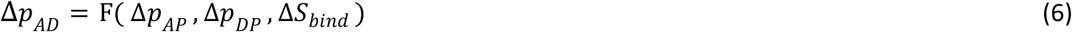

where Δ*p_AP_* = Δ*p_AP_*(*A_m_*) − Δ*p_AP_*(*A_wt_*) is the difference in amyloid-promoting propensities, Δ*p_DP_* = Δ*p_DP_*(*A_m_*) − Δ*p_DP_*(*A_wt_*) is the difference in droplet-promoting propensities, and Δ*S_bind_* = *S_bind_*(*A_m_*) − *S_bind_*(*A_wt_*) is the difference in binding mode entropy between the mutated (A_m_) and wild-type (A_wt_) residues. To define the function F in Eq. (6), we trained a support vector machine (SVM) model to fit the aggregation rates derived from experimental data. We used the recovery rates (RR) of FUS mutations [56]: ALS-associated G156E, and non-ALS mutations G156D, G156Q, G156E, G156P and defined aggregation rates as 1-RR. We thus obtained a Pearson correlation coefficient r=0.943, p=0.016 (**Figure 5A**). Then we applied this SVM model on the predicted the Δ*p_AP_*, Δ*p_DP_*, Δ*S_bind_* values of ALS-associated mutations of FUS, hnRNPA1, hnRNPA2, tau, TDP-43, TIA1 and UBQLN2 proteins (**Table S4**).

## Acknowledgements

M.F. acknowledges the financial support of HAS-11015 and GINOP-2.3.2-15-2016-00044.

## Supplementary Information

**Table S1. Datasets of droplet-driver, droplet-client, droplet-amyloid, amyloid, and yeast prion proteins and datasets of droplet-promoting and amyloid-promoting regions.**

*Columns for droplet-driver, droplet-client, droplet-amyloid, amyloid, and yeast prion proteins:* UniProt ID (relevant isoforms listed), UniProt entry name, protein name, gene names, length, organism, subcellular location, function, involvement in disease.

*Columns for droplet-promoting regions:* UniProt ID (relevant isoforms listed), UniProt entry name, protein name, length, region, organism, subcellular location, function, involvement in disease, PDB structure codes. Driver and client roles are indicated when information is available. *Columns for amyloid-promoting regions*: UniProt ID (relevant isoforms listed), function category (pathogenic, functional, not known), core region, amyloid region, UniProt entry name, protein name, length, organism, subcellular location, function, involvement in disease, PDB structure codes.

**Table S2. Prediction of droplet-promoting and amyloid-promoting regions in droplet-promoting and amyloid-promoting regions; and in droplet-driver, droplet-client, amyloid and droplet-amyloid proteins.** The number and positions of short (≥ 10 residues) and long (≥ 25 residues) droplet-promoting regions are shown. Amyloid-promoting regions (≥ residues) were identified using CamSol [35] and Tango [20].

**Table S3. Prediction of droplet-promoting and amyloid-promoting regions in the human, yeast and worm proteomes.** The number and positions of short (≥ 10 residues) and long (≥ 25 residues) droplet regions are shown. Amyloid-promoting regions regions (≥ 6 residues) were identified using Tango [20].

**Table S4. Single mutations with experimental evidence for promoting aggregation in the droplet state.** We considered only mutations that have been directly related to aggregation and not through regulatory proteins, RNA interactions, misfolding or other indirect effects.

**Figure S1.**
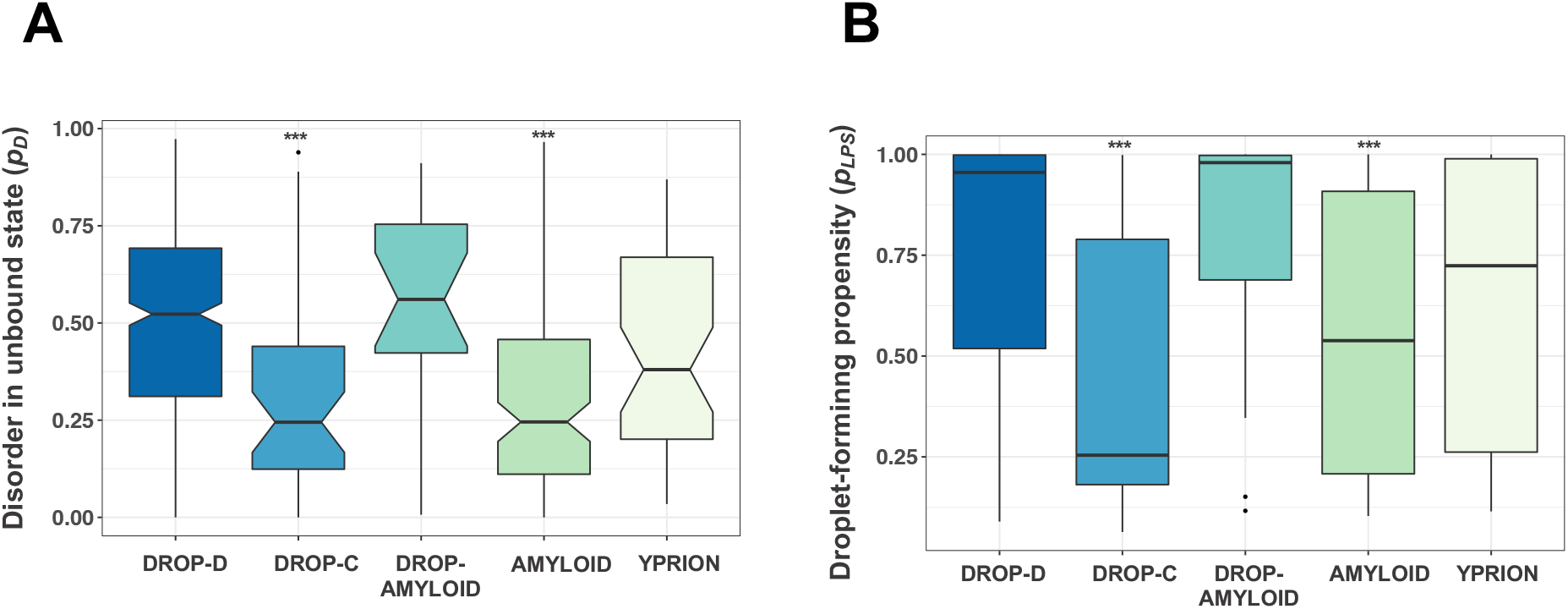
Free-state conformational entropy and droplet-forming propensities of amyloid- and droplet-forming proteins. **(A)** The free-state disorder propensity (*p_D_*) was estimated as the fraction of disordered residues by Espritz NMR [65]. Droplet-driver proteins (DROP-D, blue, **Table S1**), which spontaneously undergo liquid-liquid phase separation [27] and droplet-amyloid proteins (DROP-AMYLOID, aquamarine, **Table S1**), which exhibit both droplet and amyloid behaviors are the most disordered in the free state. Amyloid-forming (AMYLOID, green [32]) and droplet-client proteins (DROP-C, light blue, **Table S1**) are the most ordered in the free state, while yeast prions (YPRION, light green), with inheritable traits in *Saccharomyces cerevisiae* [34] have intermediate disorder values between the amyloid and droplet states. Statistical significances by Mann-Whitney test as compared to droplet-driver proteins (*** p< 10^-3^) demonstrate that amyloid-forming and droplet-client proteins are more ordered than droplet-forming proteins, while yeast prions and droplet-amyloid proteins have comparable degree of disorder to droplet-forming proteins. **(B)** The droplet-promoting propensity (p_LLPS_) was estimated by FuzDrop [27]. Droplet-driver (blue) and droplet-amyloid (aquamarine) are predicted to form droplets, whereas droplet-promoting propensities of amyloid-forming (green) and yeast prion (light green) proteins vary in a wide range. Most droplet-client proteins (light blue) are predicted not to form droplets spontaneously. Statistical significances by Mann-Whitney test as compared to droplet-driver proteins (*** p< 10^-3^) demonstrate that amyloid-forming and droplet-client proteins are less prone to form liquid droplets. Statistical significances were computed by the R program.

**Figure S2.**
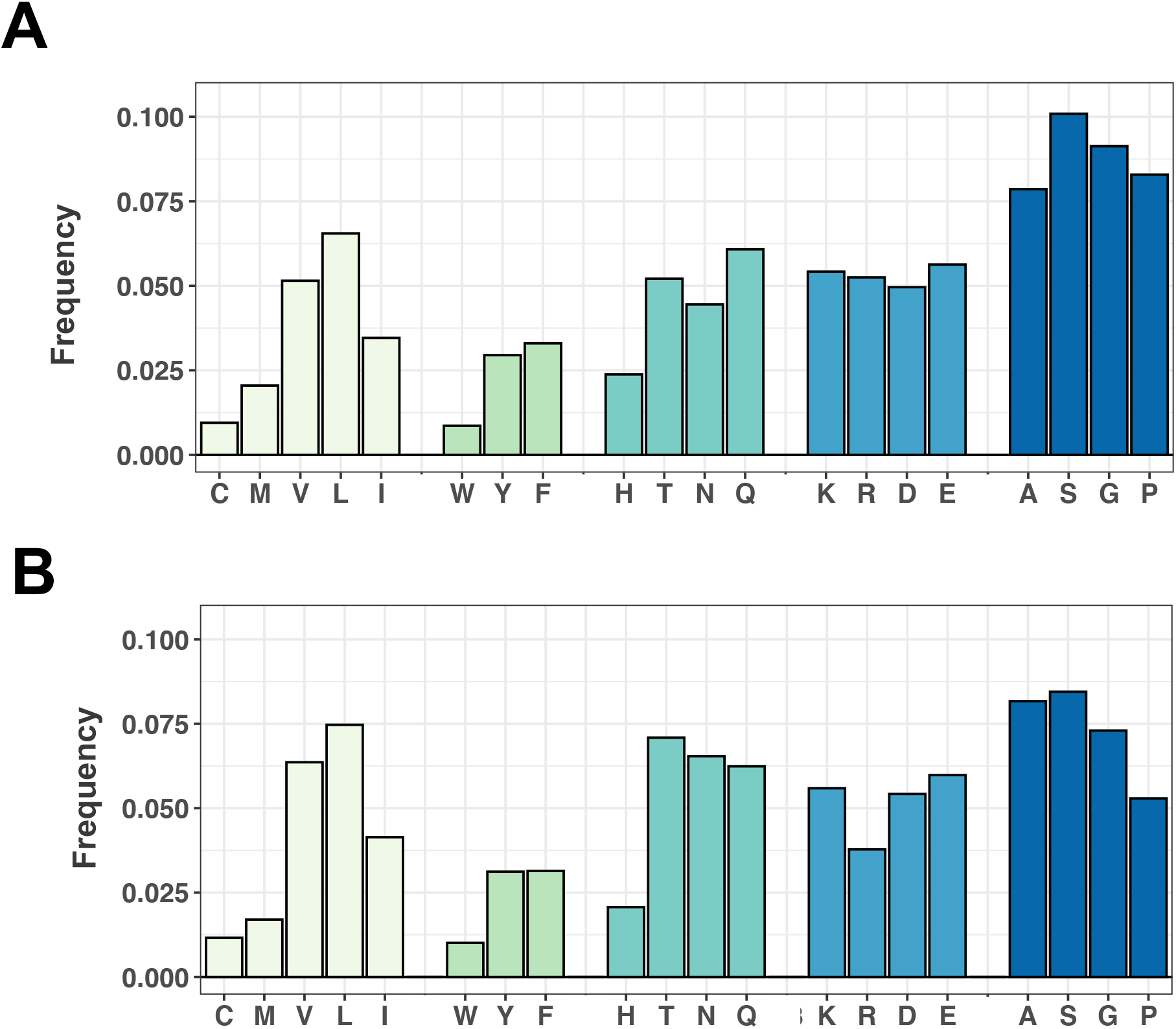
Amino acid composition of droplet-promoting (A) and amyloid-promoting (B) regions. Amino acids grouped as hydrophobic (light green), aromatic (green), hydrophilic (turquoise), charged (steel blue), disorder-promoting (dark blue). Droplet-promoting regions are enriched in disorder-promoting residues (dark blue, [43]) as compared to amyloid-promoting regions, in particular in glycine and proline in accord with previous results [44].

**Figure S3.**
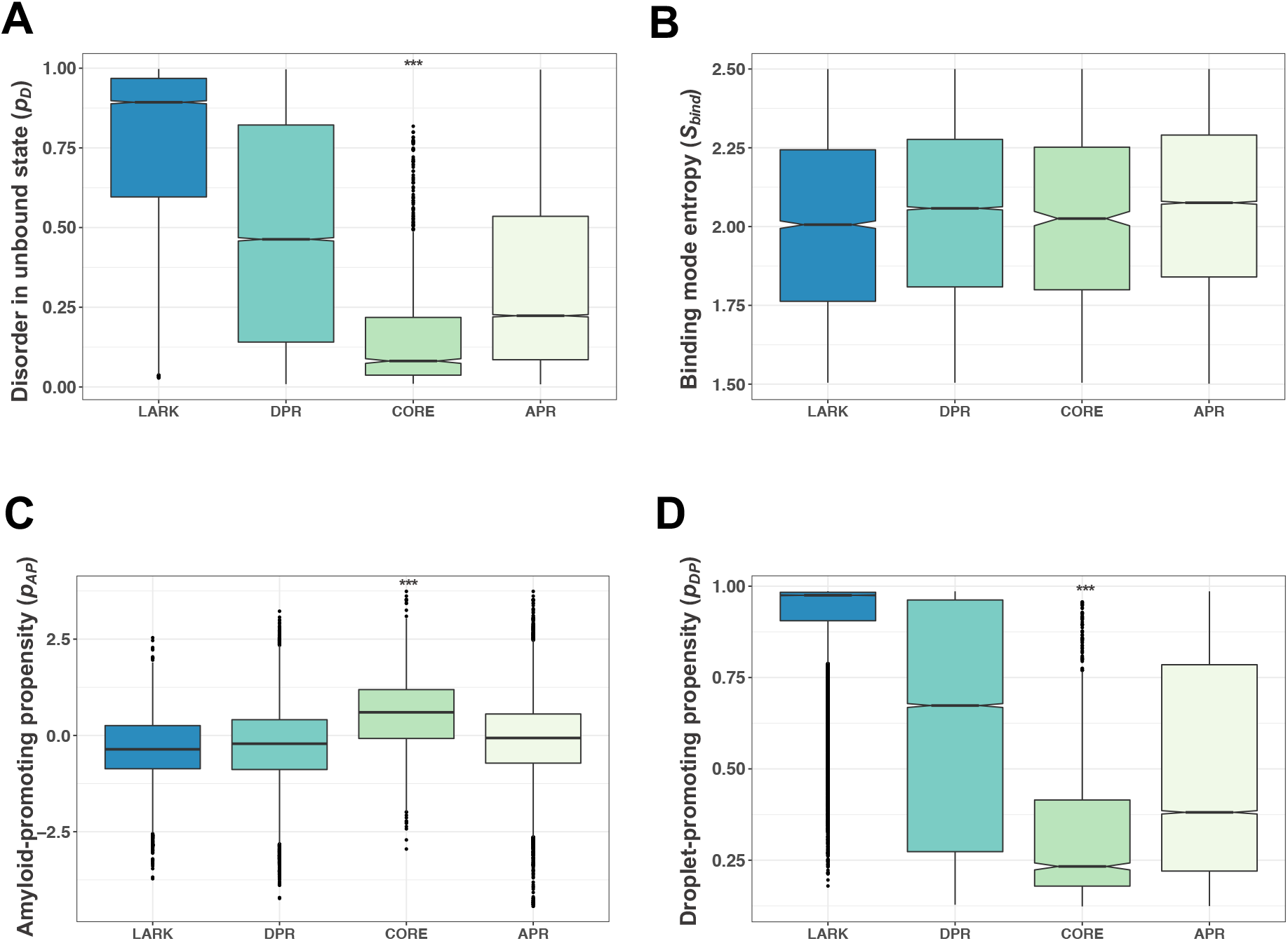
Analysis of the conformational and binding propensities of droplet-promoting regions (DPR, aquamarine) and amyloid-promoting regions (APR, light green). **(A)** Disorder propensity in the free state (*p_D_*). This propensity was estimated by the fraction of disordered residues by Espritz NMR [65]. Both droplet-promoting and amyloid-promoting regions sample a broad range of conformational states. Droplet-promoting regions are less disordered in the free state than LARKs [25] (blue), while amyloid-promoting regions are less ordered than amyloid cores (green). **(B)** Binding mode entropy (S_bind_). This property was defined by the binding mode entropy (S_bind_) as computed by FuzPred [41] (Methods, Eq. 3). Both droplet-promoting and amyloid-promoting regions exhibit larger variability of binding modes than LARKs and amyloid cores, indicating a possible conversion between ordered and disordered binding modes. **(C)** Amyloid-promoting propensity (p_AP_). This propensity was derived from the solubility scores by CamSol [35]. Droplet-promoting and amyloid-promoting regions have similar aggregation propensities, which is comparable to LARKs. Amyloid cores have lower p_AP_ values and more prone to aggregate. **(D)** Droplet-promoting propensity (p_DP_). This propensity was calculated by FuzDrop [27]. This propensity shows a sharp contrast between LARKs and amyloid-cores, while spontaneous droplet-forming propensities of droplet-promoting and amyloid-promoting regions considerably overlap, indicating that many amyloid-promoting regions also tend to phase separate. Statistical significances by Mann-Whitney test as compared to LARKs (*** p< 10^-3^) demonstrate that while amyloid cores deviate from labile beta elements, amyloid-promoting regions and droplet-promoting regions have comparable properties.

**Figure S4.**
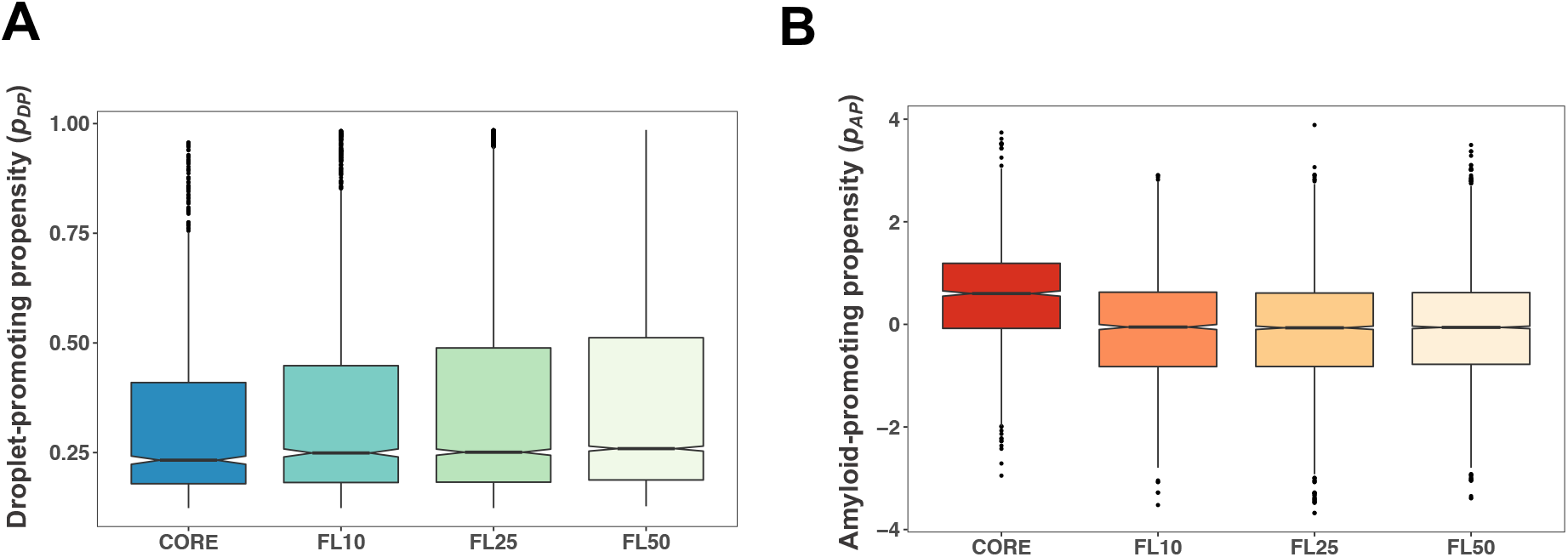
Comparison of the droplet-promoting propensity (p_DP_) and amyloid-promoting propensity (p_AP_) of amyloid cores and their flanking regions. (A,B) p_DP_ and p_AP_ as a function of the length of the flanking region. p_DP_ and p_AP_ values were evaluated in 10-residue (FL10), 25-residue (FL25) and 50-residue (FL50) flanking regions. The droplet-promoting propensity increases with the size of the flanking region. By contrast, the amyloid-promoting propensity does not change considerably with the size of the flanking region. Amyloid-promoting propensities were derived from the CamSol solubility scores [35], and droplet-promoting propensities were computed by FuzDrop [27].

**Figure S5.**
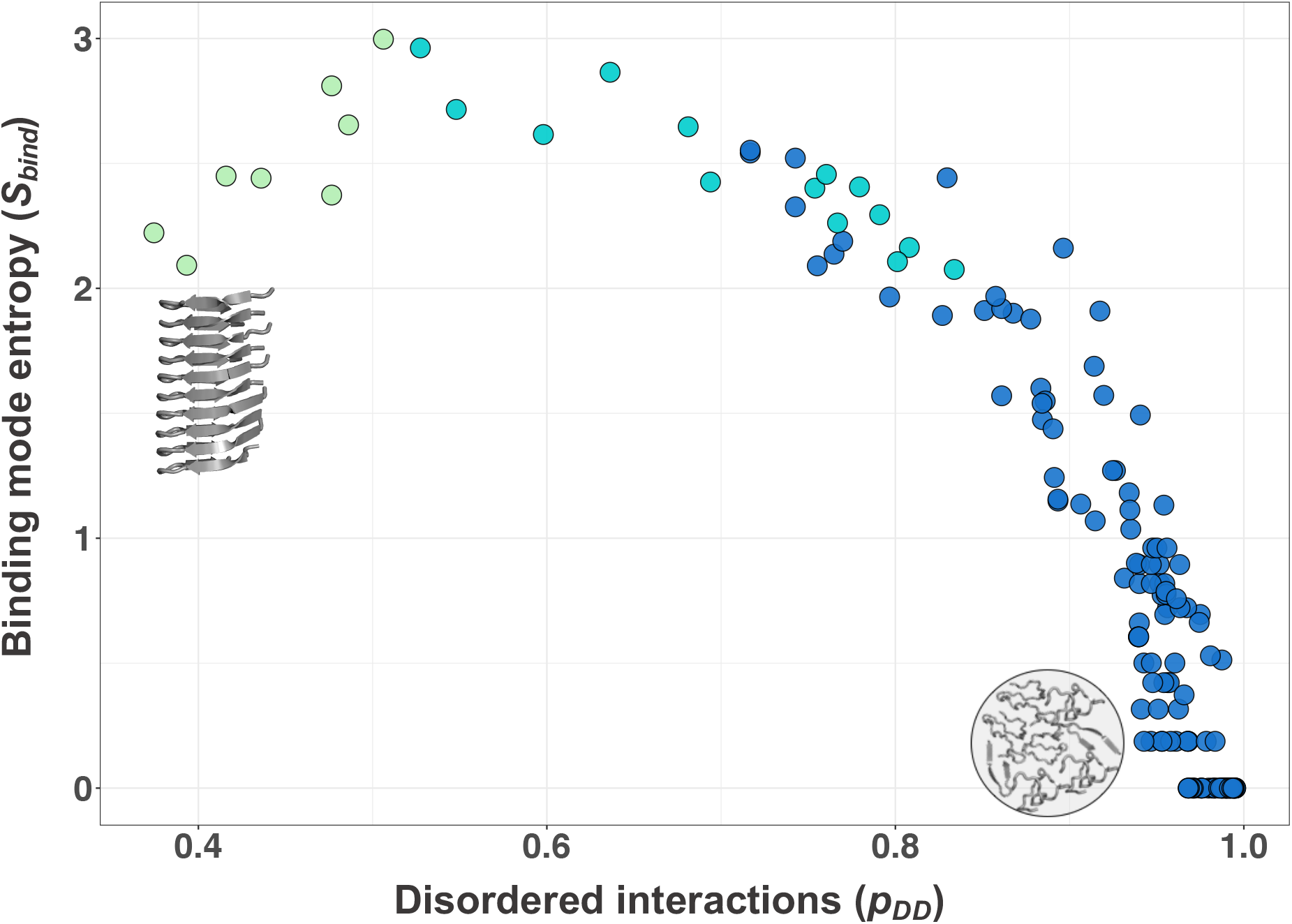
Binding landscape of the TDP-43 prion-like domain. The propensity of disordered binding modes (*p_DD_*, Methods eq. 1) by FuzPred [42] is shown on the x-axis, and the binding mode entropy (S_bind_, Methods Eq. 3), reflecting the multiplicity of interaction modes [41], on the y axis. Most residues of the TDP-43 prion-like domain sample disordered binding modes in the droplet-state (high *p_DD_*) with low variability in their interaction behavior (low S_bind_). The amyloid core region (residues 321-330, light green) exhibits more ordered binding modes with high S_bind_ values suggesting that it can sample both disordered and ordered interactions. The hot-spot region (aquamarine) also exhibits increased multimodal interactions indicating an increased likelihood of conversion to the ordered interactions.

**Figure S6.**
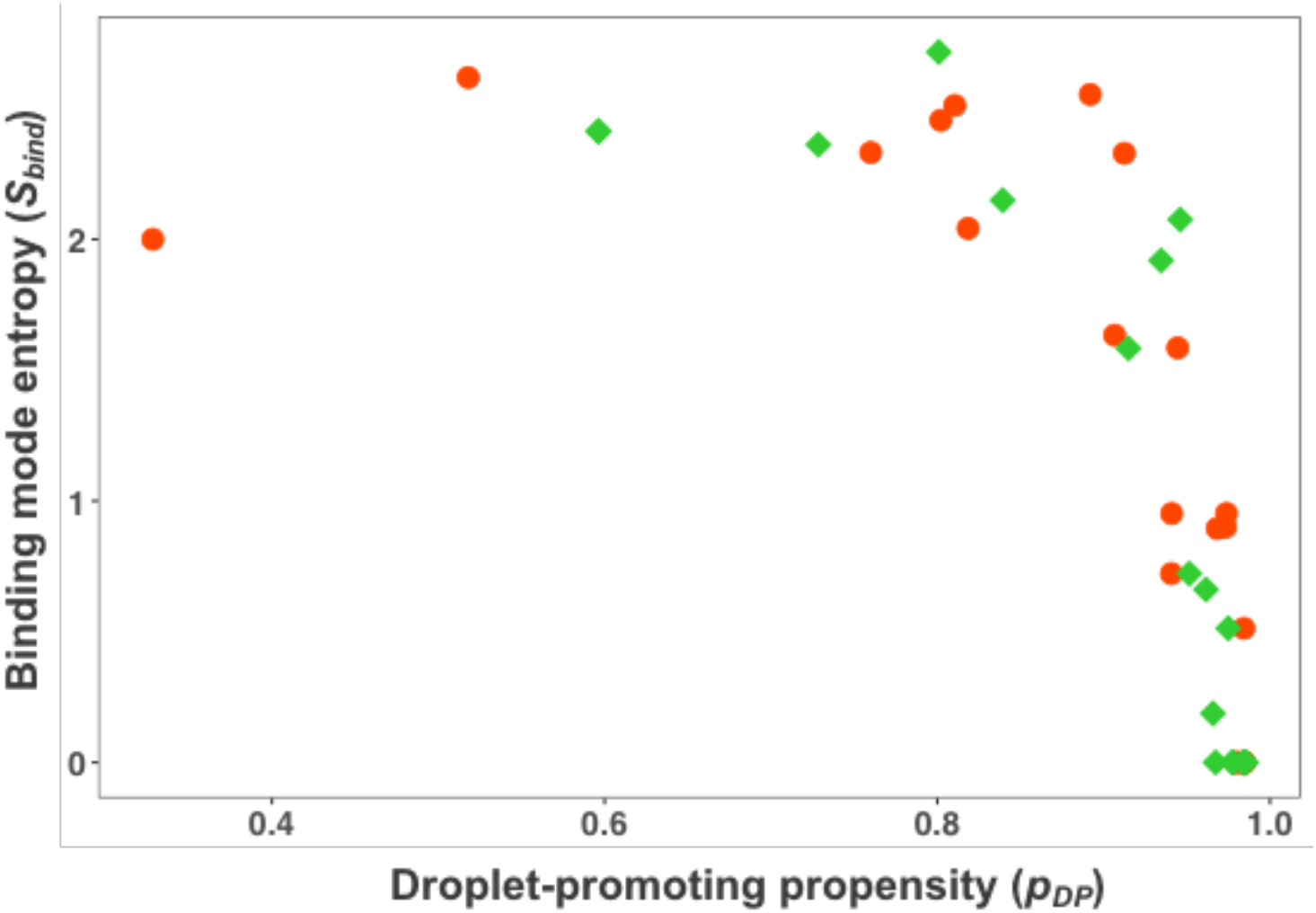
Change in the droplet landscape due to ALS-associated mutations of FUS, hnRNPA1, hnRNPA2, tau, TDP-43, TIA1 and UBQLN2. As compared to wild-type proteins (green diamond) single mutations (red circles), which were shown to promote aggregation (FUS G156E [6], G187S, G225V, G230C, G399V [48], P525L [15]; hnRNPA1 D262V [5], D214V [49]; hnRNPA2 D290V [50]; tau P301L, P301S, A152T [51]; TDP-43 A321V [47], G298S, M337V [52], A315T [53]; TIA1 P262L, A381T, E384K [54]; UBQL2 P506T [55]) (**Table S4**) shift the landscape towards lower droplet-promoting propensity (p_DP_) [27] and higher binding mode entropy (S_bind_) [41].

